# An agnostic study of associations between ABO and RhD blood group and phenome-wide disease risk

**DOI:** 10.1101/2021.01.28.428569

**Authors:** Torsten Dahlén, Mark Clements, Jingcheng Zhao, Martin L Olsson, Gustaf Edgren

## Abstract

There are multiple known associations between the ABO and RhD blood groups and disease. However, no systematic population-based studies elucidating associations between a large number of disease states and blood group have been conducted. Using SCANDAT3-S, a comprehensive nationwide blood donation-transfusion database, we modelled outcomes for 1,217 disease categories including 70 million person-years of follow-up, accruing from 5.1 million unique individuals. We discovered 49 and 1 associations between a disease and ABO and RhD blood group, respectively, after adjustment for multiple testing. We identified new associations such as kidney stones and blood group B as compared to O. We also expanded previous knowledge on other associations such as pregnancy-induced hypertension and blood group A and AB as compared to O and RhD positive as compared to negative. Our findings generate strong further support for previously known associations, but also indicate new interesting relations.

## Introduction

The blood group antigens of the ABO and RhD systems play a pivotal role in transfusion medicine because of their role in the safe administration of blood transfusions. In addition, these cell surface antigens have been demonstrated to have direct effects on the susceptibility for several diseases (1–3). One of the first such studies was published in 1962, demonstrating a relation between the ABO system and ischemic heart disease (4). Multiple subsequent studies have revealed associations with a range of diseases, with a prominent example being a decreased risk of thromboembolic events and increased risks of some hemorrhagic events in individuals with blood group O (1, 5, 6). The difference in thrombotic and hemorrhagic phenotypes has been attributed to variability in levels of *Factor VIII* and *von Willebrand factor* (vWF), where ABO status may explain as much as 30% of this variability (1, 7) (8–10) Other prominent examples include associations with risks of a number of infectious diseases, to the extent that the allele distribution of the blood group antigens has evolved to reflect some areas endemic to these infectious diseases (11). This is in part true for infectious disease such as *Plasmodium falciparum* malaria, *Helicobacter pylori* and *Vibrio cholera*, where ABO blood groups are involved in different aspects of pathogenesis, from microbe attachment and entry into cells to subsequent disease development and severity of disease *(2, 11–13).* The RhD antigen, on the other hand, has a less clear link to health outcomes. RhD status has mainly been linked to alloimmunization of the pregnant women with hemolytic disease of the fetus and newborn (HDFN) as a consequence (14). Beyond these direct effects, little is known about its role in disease pathogenesis. The difference between RhD positive and negative blood group, is the presence or absence of the RhD protein on the red blood cell surface. However, both individuals with and without RhD possess the homologous RhCE protein and Rh-associated glycoprotein (RhAG) on their red cells. Thus, functions carried out by RhD are likely performed RhCE and RhAG in RhD-negative individuals, and this redundancy may in part explain the scarcity of findings related to RhD status (15).

Using the Scandinavian Donation and Transfusion (SCANDAT) database, we have previously studied associations between ABO blood groups and cancer subtypes, cardiovascular and thromboembolic disease, the occurrence of dementia and degradation of bioprosthetic aortic valves in relation to ABO blood group (6, 16-18). However, these and most other prior studies into the association between ABO blood group and disease outcomes have been limited by potentially misdirected *a priori* hypotheses and phenome-wide disease associations have not been thoroughly explored in a systematic manner. Therefore, in the current study, we aimed to agnostically investigate the association between ABO and RhD blood group and disease occurrence for a large number of disease phenotypes using large-scale population-based Swedish healthcare registries.

## Results

Characteristics of the main and validation cohorts are presented in Table 1. When combining the main and validation cohort, there were a total of 5.1 million unique individuals. The main cohort consisted of 4.2 million individuals who at any point had undertaken an ABO and RhD blood antigen test. The distribution of A, AB, B, O were 47%, 5%, 10% and 38%, respectively, and 84% of individuals were RhD positive. Women constituted 60% of the cohort. The median age at cohort entry was 52 years (interquartile range [IQR], 30-71) and the median year of birth was 1949 (IQR, 1931-1971).

**Table 1.** Baseline characteristics of main and validation cohort

|  | Main cohort |  | Validation cohort |  |
| --- | --- | --- | --- | --- |
| **Number** | 4 204 234 |  | 1 197 522 |  |
| **Age, median (IQR)** | 52 | (30-71) | 30 | (23-41) |
| **Year of birth, median (IQR)** | 1949 | (1931-1971) | 1966 | (1953-1978) |
| **Sex, %** | 60 |  | 49 |  |
| **Blood group, %** |  |  |  |  |
| A | 47 |  | 45 |  |
| AB | 5 |  | 5 |  |
| B | 10 |  | 11 |  |
| O | 38 |  | 39 |  |
| **RhD positive, %** | 84 |  | 82 |  |

IQR: interquartile range.

Not accounting for censoring due to disease events, the main cohort accrued a total of 49.9 million person-years of follow-up, 23.7 million in blood group A, 2.3 million in blood group AB, 4.9 million in blood group B, and 18.9 million in blood group O.

Of the original 1,217 disease categories, 1,090 remained available for analyses after excluding disease categories with fewer than 50 events. The median number of events per disease category in the main cohort was 4,748 (IQR, 869-231,166). A meta-summary of results of regression analyses is presented in Table 2, and graphically in the form of a volcano plot in Figure 1 (also, as an interactive, online variant as Supplementary Figure 1). Alternatively, results are also presented as an ICD chapter-based, variant Manhattan plot in Figure 2 (also as an interactive, online variant as Supplementary Figure 2). Overall, in the main cohort and before FDR adjustment for multiple testing, there were 343 and 98 statistically significant associations for the ABO and RhD blood group systems and unique disease categories, respectively. Of these, a total of 143 (41%) and 13 (13%) associations between blood group and unique outcome remained statistically significant for ABO and RhD blood group systems, respectively, after FDR adjustment. For the ABO system, IRRs for statistically significant associations after FDR-adjustment ranged from 0.57 to 0.99 for negative associations, and from 1.01 to 1.52 for positive associations. For RhD status, IRRs ranged from 0.90 to 0.97 for negative associations, and from 1.02 to 1.08 for positive associations. Details of all associations identified after FDR are presented in Supplementary Table 3 (for ABO blood groups) and Supplementary Table 4 (for RhD).

**Table 2.** Meta-summary of results

|  | Main cohort |  |  |  |  | Validation cohort |  |  |  |  |
| --- | --- | --- | --- | --- | --- | --- | --- | --- | --- | --- |
| **Individuals** | 4 204 234 |  |  |  |  | 1 197 522 |  |  |  |  |
| **Person-years, sum** | 50M |  |  |  |  | 22M |  |  |  |  |
| **Events, median (IQR)** | 4748 (869-23166) |  |  |  |  | 7129 (2464-19973) |  |  |  |  |
|  | A | AB | B | ABO Total | RhD positive | A | AB | B | ABO Total | RhD positive |
| **Before adjustment** | 229 | 129 | 179 | 537 | 106 | 66 | 37 | 44 | 147 | 6 |
| **Positive effects, N** | 150 | 79 | 107 | 336 | 61 | 48 | 23 | 22 | 93 | 3 |
| **Negative effects, N** | 79 | 50 | 72 | 201 | 45 | 18 | 14 | 22 | 54 | 3 |
| **After adjustment\*** | 108 | 56 | 70 | 234 | 13 | 26 | 13 | 10 | 49 | 1 |
| **Positive effects, N** | 66 | 38 | 36 | 140 | 10 | 19 | 10 | 6 | 35 | 1 |
| **Negative effects, N** | 42 | 18 | 34 | 94 | 3 | 7 | 3 | 4 | 14 | 0 |
| **Positive IRR, median (range)** | 1.05(1.01-1.72) | 1.09(1.03-1.52) | 1.09(1.02-1.39) |  | 1.05(1.02-1.08) | 1.1(1.03-1.57) | 1.32(1.07-1.89) | 1.5(1.07-1.64) |  | 1.12(1.12-1.12) |
| **Negative IRR, median (range)** | 0.95(0.77-0.99) | 0.92(0.74-0.97) | 0.92(0.57-0.98) |  | 0.97(0.9-0.97) | 0.92(0.86-0.95) | 0.84(0.81-0.87) | 0.88(0.83-0.93) |  | - |

\* In the main and validation cohort FDR and Bonferroni adjustment was conducted, respectively.

**Figure 1.**
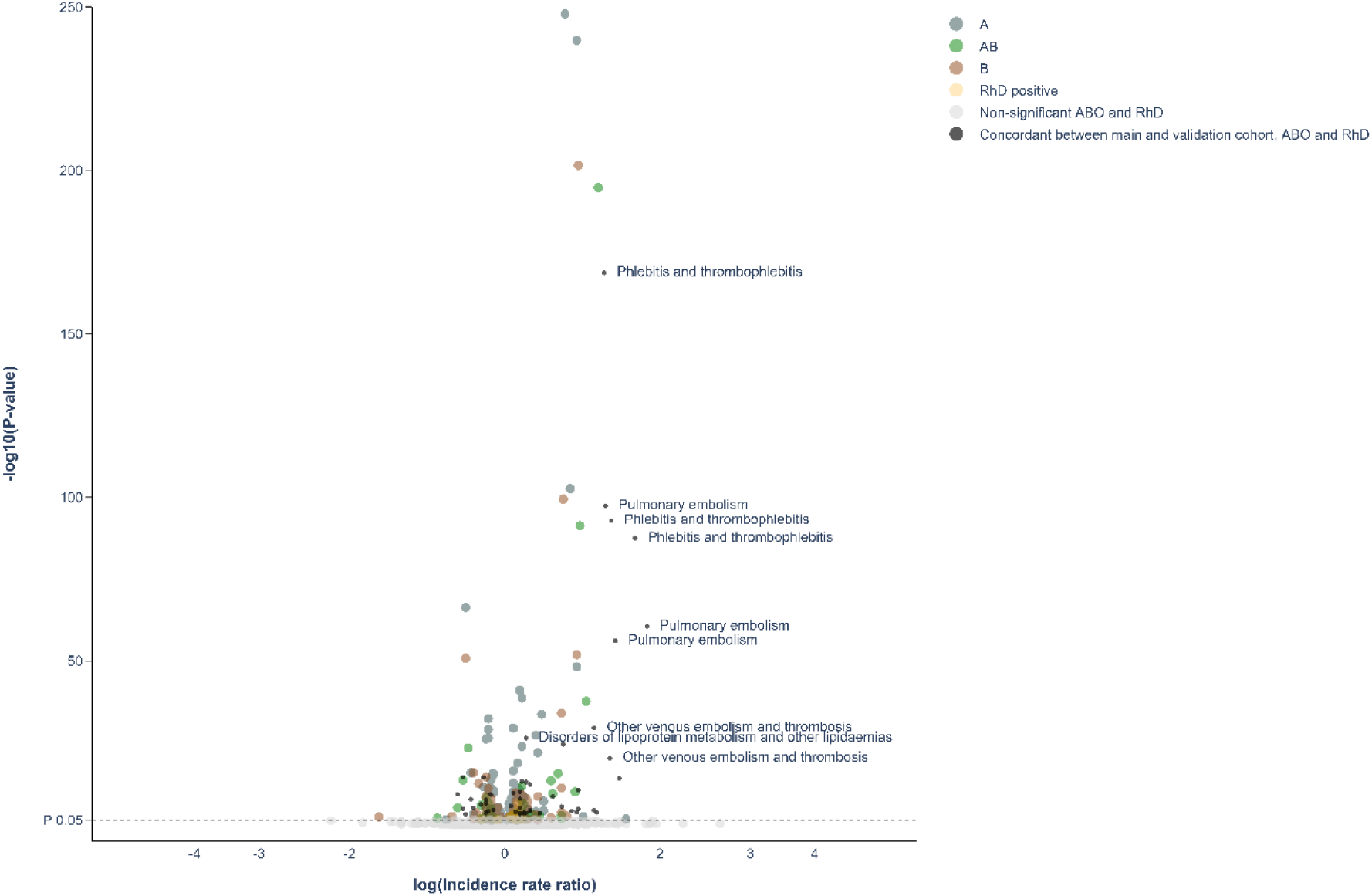
Volcano-plot of all findings from main and validation cohort Volcano-plot depicting spread of P values of significant and non-significant ABO and RhD blood groups for main cohort and validated results (live version available as online Supplementary Fig 1). The labels represent the 9 findings with the lowest P-value in the validation cohort.

**Figure 2.**
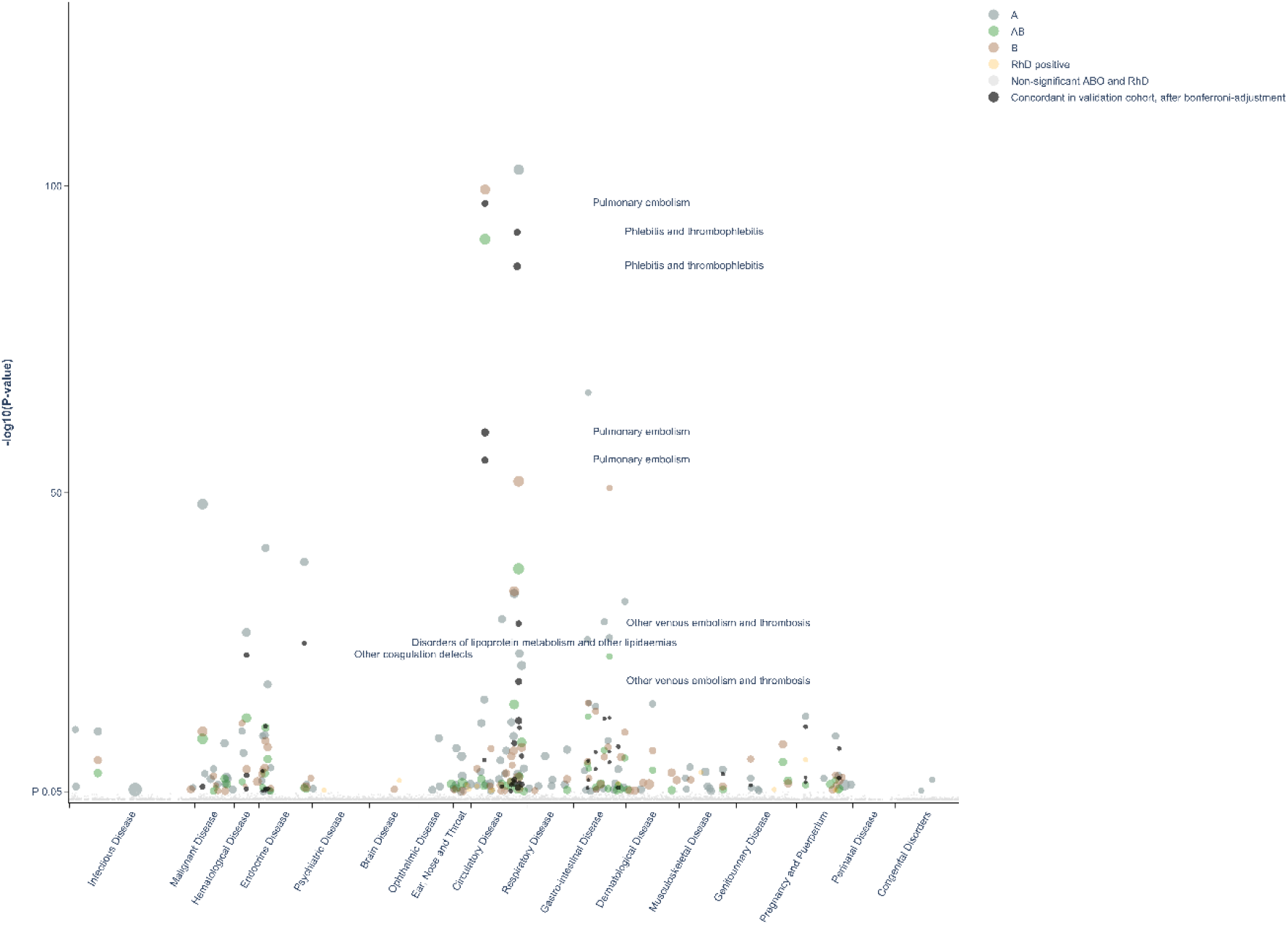
Manhattan-plot of all findings from main and validation cohort mapped by ICD chapter. Manhattan-plot depicting distribution of P-values for significant and non-significant associations between ABO and RhD blood groups and available outcomes for main cohort and validated results mapped by disease chapter in ICD (live version available as online Supplementary Fig 2).

**Figure 3.**
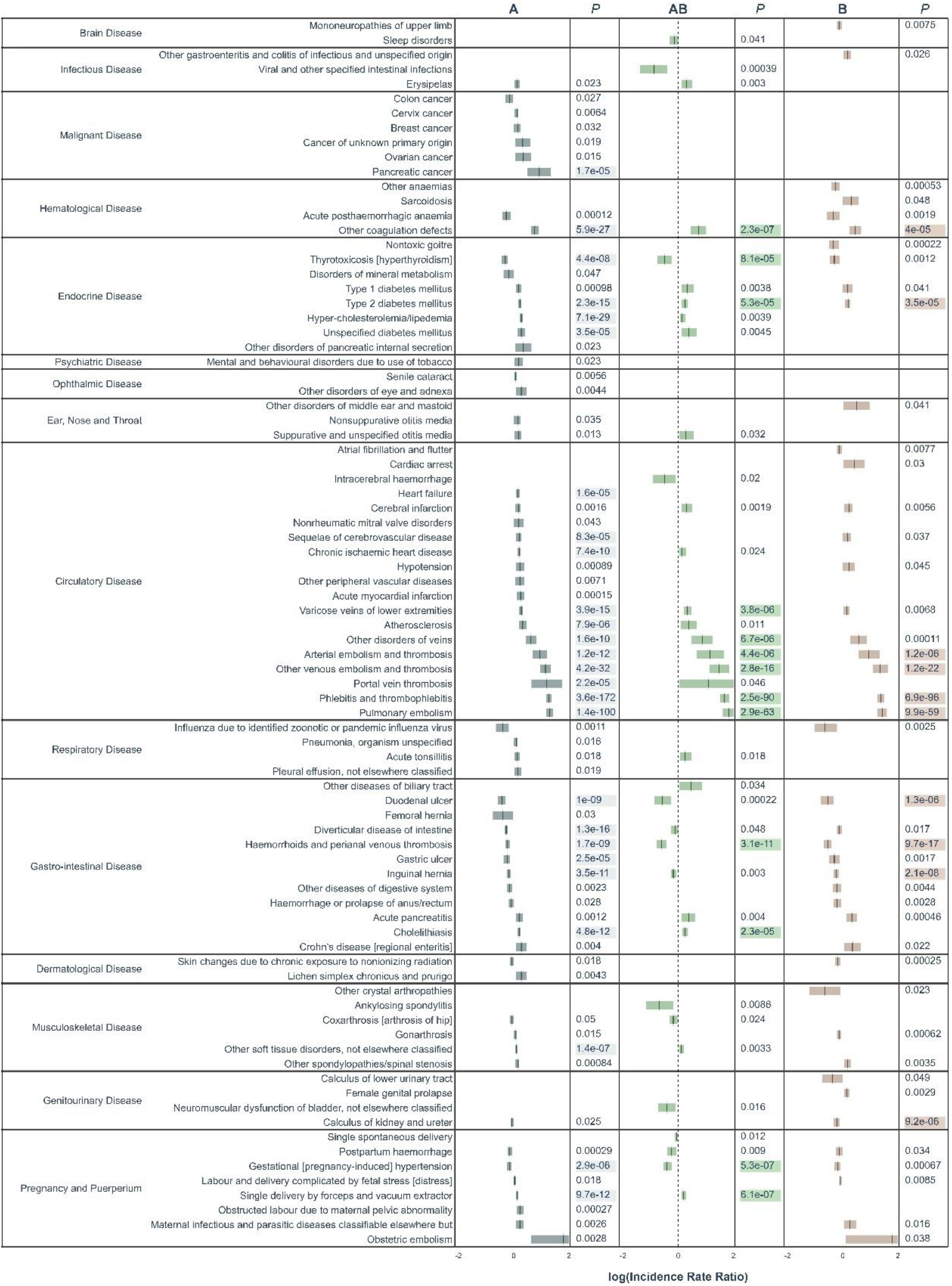
All significant un-adjusted findings from the validation cohort. Significant disease categories in the validation cohort. Blood group as compared to blood group O and log10(IRR) displayed with 95% confidence bands. All P values are raw, highlighted P value indicates associations that remained statistically significant also after Bonferroni-adjustment.

In our validation cohort, consisting of almost 1.2 million blood donors accruing 22 million person-years of follow-up, we validated the findings from the significant disease categories from the first analysis. Among the 143 and 13 significant disease categories for ABO and RhD, respectively, the median number of events was 7,129 (interquartile range 2,464-19,973). Before multiple testing adjustment, we identified 160 associations between a blood group in 147 and 6 disease categories, for the ABO and RhD blood group, respectively. After Bonferroni-adjustment, there were 49 and 1 associations remaining between ABO and RhD blood group, respectively (Table 2).

A number of previously well-established associations were seen among the Bonferroni-adjusted results. For thrombosis, blood group A had a higher risk as compared to O (e.g., pulmonary embolism, IRR 1.57 [95% CI, 1.51-1.64] and portal vein thrombosis, IRR 1.51 [95% CI, 1.25-1.83]). Bleeding disorders were more frequent in blood group O as compared to A (e.g., gastric ulcer, IRR 0.92 [95% CI, 0.88-0.95] and duodenal ulcer was, IRR 0.86 [95% CI, 0.82-0.9]). Thyrotoxicosis was also less common in blood group A and AB as compared to blood group O (with IRRs of 0.90 [95% CI 0.86-0.93] and 0.84 [95% CI, 0.77-0.92], for A and AB, respectively). Pregnancy-induced hypertension was less common in blood groups A and AB, as compared to blood group O (with IRRs of 0.95 [95% CI, 0.92-0.97] and 0.87 [95% CI, 0.83-0.92] for A and AB, respectively). Pancreatic cancer was the only malignancy that remained associated with a blood group, specifically blood group A as compared to O (IRR, 1.29; 95% CI, 1.19-1.40). A new finding was that of calculus of the kidney and ureter, which were found to be less common in blood group B as compared to O (IRR 0.93 [95% CI, 0.89-0.96]). Cholelithiasis, which has been disputed, was more common in blood group A and AB as compared to blood group O (with IRRs of 1.07 [95% CI, 1.05-1.09] and 1.09 [95% CI, 1.05-1.13] for A and AB, respectively).

In disease categories not significant after Bonferroni-adjustment, some findings exhibited particularly strong effects, such as for viral and other specified intestinal infections, where blood group AB had a significantly lower risk, as compared to O (IRR 0.74; 95% CI, 0.62-0.87). There was a lower risk for ankylosing spondylitis in blood group AB as compared to O (IRR 0.79; 95% CI, 0.67-0.94), and for acute pancreatitis, again with a lower risk in blood group AB as compared to O (IRR 1.14; 95% CI, 1.04-1.24). For the RhD positive as compared to negative, only one disease category remained statistically significant after Bonferroni-adjustment, namely pregnancy-induced hypertension (IRR 1.12; 95% CI, 1.09-1.16). Strong effects identified in the main cohort but not in the validation cohort were hereditary factor VIII deficiency in blood group B (IRR, 0.57; 95% CI, 0.42-0.77), well differentiated thyroid cancer in blood groups AB (IRR 0.74; 95% CI, 0.62-0.88) and B (IRR 0.79; 95% CI, 0.70-0.90), measles in blood group A (IRR 1.72; 95% CI, 1.23-2.39), as well as both erythema nodosum (IRR 1.32; 95% CI, 1.15-1.53) and sarcoidosis in blood group B (IRR, 1.15; 95% CI, 1.08-1.23), as compared to blood group O.

## Discussion

In this large cohort study of 5.1 million unique persons followed over 70 million person-years, we performed an agnostic analysis of associations between the ABO and RhD blood groups and the risk of 1,217 distinct disease categories. After multiple testing adjustment and comparison with a validation cohort, 50 and 5 associations between disease categories and blood group for ABO and RhD remained significant, respectively. Overall, we were able to confirm a number of previously known associations such as risk of thrombosis and hemorrhagic events. In addition, we also identified novel associations, some with firm evidence and valid even after conservative Bonferroni-adjustment, in the validation cohort, including for example calculus of the kidney and ureter. Furthermore, being the largest study so far, we also found that blood group A and AB had a lower risk of gestational hypertension, compared to blood group O, which has previously been disputed (19–21). Of the identified associations, we speculate that some associations may be driven by increased screening due to other concomitant diseases that are associated with blood groups, which might be the case for hyperlipidemias that are screened for in heart disease. Most of the investigated disease or disease groups, however, do not seem to be strongly influenced by the ABO blood group of the individual.

This is hitherto the largest study investigating blood group antigens and disease occurrence in an effort to find novel and confirm previously known associations. There are some particular strengths to our approach and data. Most notably, the study was based on a very large study population, representing one third of the Swedish population, with long-term and unbiased follow-up. This ensures both the reliability and the generalizability of the results. The agnostic approach also has the advantage of not being based on specific pre-set conceptions of specific disease categories and possible associations of blood group. All disease categories are treated equally and investigated using the same principles effectively removing researcher bias. Moreover, the data has been collected prospectively in various high-quality health-care registries during a long time period with almost complete coverage. In addition, the fact that all blood group data in the SCANDAT3-S database were collected from clinical transfusion registers – the quality of which is essential for the safe administration of blood transfusions – ensures that there should be little or no errors in the blood group coding. Similarly, while the validity of the outcomes registration certainly varies between the different disease categories, the degree of such misclassification is unlikely to vary between blood groups, and so it should not affect the magnitude of point estimates.

The current study is limited by several factors. One such factor is the disease classification scheme used, based primarily on ICD revision 10 categories. Smaller disease entities were not accounted for and thus there may be true associations that were missed. It is thus possible that some of the associations that we reported were driven by multiple unknown associations within a specific disease category that may have unequal, or even detrimental, effects on the outcome. However, we believe that this limitation is an opportunity for further sub-categorized investigations in the future when even more events and follow-up time are available.

Another limitation that prevents strong casual inference is the possibility some of the observed associations between ABO blood group and disease categories were driven by other disease associations with ABO blood group. This might, for example, be the case for the associations between blood group A and diabetes as well as hyperlipidemia, which are potentially driven not by a causal association but possibly instead by an association between blood group A and ischemic heart disease, at the occurrence of which diabetes and lipid disorders are screened for and thus frequently diagnosed. We cannot exclude the possibility that some of the other associations were driven by similar non-causal mechanisms.

To limit the possibility of false positive findings we handled over-dispersed Poisson models using Quasi-Poisson and also in the main cohort applied the FDR approach, described by Benjamini and Hochberg, and then utilised a Bonferroni-adjustment on the sub-grouped outcomes in the validation cohort. The aim of this approach was to reduce type 1 errors without being overly conservative by first conducting an explorative analysis in the main cohort. We also employed a confirmatory analysis with a validation cohort to further limit the possibility of false positive findings. However, because the validation cohort was both smaller and consisted only of blood donors, who were selected for their good health, the ensuing smaller number of events may result in failure to detect potentially interesting associations. However, in the validation cohort, only approximately 1% of the categories had fewer events than 50. It may still be informative to consider also some of the associations from the main cohort that were not corroborated in the validation cohort. This is exemplified by pancreatic cancer where we saw an increased risk in blood group AB and B in the main cohort (IRR, 1.37 and 1.129, p<0.00001 and p<0.00001 for AB and B, respectively) and a similar, yet non-significant effect in the validation cohort (IRR, 1.14 and 1.26, p-value 0.8 and 0.6 for B and AB, respectively). This also expands to the non-findings in terms of cancerous disease were multiple relationships that have previously been demonstrated but not in the validation cohort after Bonferroni-adjustment. This strengthens our decision to not limit the presentation of findings to only disease categories identified in the conservative Bonferroni-adjusted analysis (17).

Still, after these limitations we believe that our findings support and generate strong further evidence for previously known associations and indicate new and interesting relationships for disease such as calculus of the kidney and ureter, pregnancy-induced hypertension, well-differentiated thyroid cancer and sarcoidosis. The new set of associations should be validated in other cohorts but also investigated using a mechanistic approach for a possible causal and biological interaction.

## Materials and Methods

### Study population and study design

Individuals in the study were identified using an updated version of the Scandinavian donations and transfusion database (SCANDAT3-S). This database includes close to 8 million individuals who have donated blood, received a blood transfusion, or have had blood group testing done for other reasons. Other reasons for blood group testing would typically be pre-emptive testing e.g., before surgery or in antenatal care. The database contains detailed information on blood donations, transfusions as well as blood group antigen and antibody testing results and is thoroughly described elsewhere (22). It is nationally complete since 1995, but information dates back to 1968 with various levels of completeness, mainly depending on the geographical region. Using unique national registration numbers assigned to all inhabitants of Sweden, the SCANDAT3-S database has been linked to a range of national health outcomes registers, for hospital care, cancer, cause of death and drug prescriptions ^19^. From SCANDAT3-S, we extracted information on ABO and RhD blood group and created a main cohort and a validation cohort. The main cohort consisted of all individuals who were born in Sweden where at least one parent was born in Sweden and who, for any reason, had undergone ABO and RhD blood group typing with a conclusive result, but who did not donate blood within 90 days of the test. Person-time for blood donors were excluded from the main cohort to maximize the representativeness of the study population. In the validation cohort, we included all individuals in the SCANDAT3-S database who had ever donated blood. As such, an individual could contribute person-time in both cohorts, such as in the case a person started to donate blood more than 90 days later from a blood grouping test that was initially performed for other reasons. The person-time before blood donation would contribute to the main cohort censoring at entry in the validation cohort starting at the time of blood grouping before the blood donation.

### Outcomes

We defined and studied a large number of disease categories. Non-cancer disease categories were based on discharge diagnoses from the national patient register, which covers all hospital inpatient care in Sweden since 1987 and all specialist outpatient care since 1997, and from the Cause of Death register, which records underlying causes of death for all persons in Sweden since 1964 (23, 24). Because the 10th revision of the International Classification of Disease (ICD) was implemented in 1997, we limited outcomes ascertainment to events from 1997 or later to avoid inconsistencies between ICD revisions. Cancer outcomes were based on the Cancer Register, which records all incident cancer cases in Sweden since 1958 (25). All of these registries are held and maintained by the Swedish National Board of Health and Welfare and have a high level of completeness and accuracy. Dates of death and emigration were obtained from population registers kept by Statistics Sweden.

Details of non-cancerous disease categories are presented in Supplementary Table 1. Non-cancer diseases were classified into disease categories based on the first 3 codes of the diagnosis, according to the ICD-10 codebook. We did not consider external causes of disease, traumatic injuries or symptom-based codes as these were deemed unlikely to be related to blood group antigens. Cancer disease categories were based on anatomical coding using the 7th revision of the ICD for all non-hematological malignancies and the 8th revision of the ICD for hematological malignancies. For details of cancer categories, see Supplementary Table 2 (SAS code for cancer disease grouping is available upon request).

In total we considered 1,217 distinct disease categories. After database construction we excluded disease categories with fewer than 50 events before analysis as we would be unlikely to detect sufficient events in the validation cohort in categories with fewer than 50 events in the main cohort.

### Statistical methods

All persons were followed from the date of the first blood grouping test, from their 18th birthday, or from January 1^st^, 1997, whichever occurred last. Follow-up was extended until the first incident event in each disease category, emigration, death or December 31^st^, 2017, whichever occurred first. A person could thus be included in follow-up for all disease categories investigated.

Descriptive statistics were presented for cohort baseline data. For the main analysis we used a Poisson regression model. In the model we incorporated the following covariates: ABO blood group (A, AB, B or O), RhD status (weak or category expression variants were excluded), sex, calendar-period, and age. A restricted cubic spline functions with 4 or 5 knots placed according to Harrell’s method were applied to the age and calendar period covariates (26). The regression model was fitted separately to each disease category resulting in incidence rate ratios (IRR) for each ABO blood group and RhD-status using blood group O and RhD negative as reference, respectively. Wald’s method was used to construct 95% confidence intervals. Equi-dispersion was tested using a Lagrange multiplier test. For disease categories where data demonstrated significant over- or under-dispersion after also performing the same analysis but reducing the number of knots from 5 to 4, analyses were instead run using quasi-Poisson regression.

Multiple testing was handled using a two-stage approach. First, in the exploratory analysis using the main cohort, we applied a false discovery rate (FDR) adjustment of raw p-values assuming positive dependency of stochastic ordering between outcomes. Second, in the confirmatory analysis using the validation cohort, we used the disease categories with significant effects from explorative analysis, with results presented both without adjustment and using a Bonferroni-adjustment. In effect, this allowed us to limit type 1 errors presenting confirmed associations with high certainty, but still not to compromise type 2 errors for future confirmatory analysis in other cohorts.

## Ethical approval

This study has been approved by the regional Stockholm County Board of Ethics Committee (ref nr: 2018/167-31).

## Acknowledgments

The creation of the SCANDAT3-S database and the conduct of this study was made possible by a grant to Dr Edgren from Swedish Research Council (2017–01954). Dr Edgren is supported by Region Stockholm (clinical research appointment). Dr Dahlén is supported by Region Stockholm (clinical research appointment). Dr Zhao is supported by the Clinical Scientist Training Program and the Research Internship Program, both at Karolinska Institutet.

## Supplementary Files

**Live figures S1 to S2.** Volcano, representing Fig. 1 in the main manuscripts but with live features, (S1) and manhattan-plot, representing Fig. 2 in the main manuscripts but with live features (S2), standalone html file with integrated javascript libraries.

**Table S1.** Non-cancer disease categories with ICD codes and names of categories, searchable html-file.

**Table S2.** Cancer disease categories, searchable html-file.

**Table S3.** All significant results from ABO analysis in the main cohort after FDR adjustment, searchable html-file.

**Table S4.** All significant results from RhD analysis in the main cohort after FDR adjustment, searchable html-file.

**Table S5.** All findings with P-value less than 0.05 in the validation cohort in the ABO analysis, Bonferroni robust findings labeled, searchable html-file.

**Table S6.** All findings with P-value less than 0.05 in the validation cohort in the RhD analysis, Bonferroni robust findings labeled, searchable html-file.

